# Single intravitreal administration of a tetravalent siRNA exhibits robust and efficient gene silencing in rodent and swine photoreceptors

**DOI:** 10.1101/2023.09.20.558641

**Authors:** Shun-Yun Cheng, Jillian Caiazzi, Annabelle Biscans, Julia F. Alterman, Dimas Echeverria, Nicholas McHugh, Matthew Hassler, Samson Jolly, Delaney Giguere, Joris Cipi, Anastasia Khvorova, Claudio Punzo

## Abstract

Inherited retinal dystrophies caused by dominant mutations in photoreceptor-expressed genes, are a major cause of irreversible vision loss. Oligonucleotide therapy has been of interest in diseases that conventional medicine cannot target. In the early days, small interfering RNAs (siRNAs) were explored in clinical trials for retinal disorders with limited success due to a lack of stability and efficient cellular delivery. Thus, an unmet need exists to identify siRNA chemistry that targets photoreceptor-expressed genes. Here we evaluated 12 different fully chemically modified siRNA configurations, where the valency and conjugate structure were systematically altered. The impact on retinal distribution following intravitreal delivery was examined. We found that the increase in valency (tetravalent siRNA) supports the best photoreceptor accumulation. A single intravitreal administration induces multi-months efficacy in rodent and porcine retinas while showing a good safety profile. The data suggest that this configuration can treat retinal diseases caused by photoreceptor-expressed genes with 1-2 intravitreal injections per year.

## Introduction

RNA interference is an endogenous molecular mechanism that utilizes small noncoding RNAs to regulate gene expression by targeting the complementary mRNA for degradation^1^. The discovery of small interfering RNAs (siRNA) in gene regulation advanced the prospect of pharmacological approaches to treat diseases that conventional drug therapies cannot achieve^2^. During the early stages of siRNA therapy development, most oligonucleotide designs used naïve and partially modified configurations^3^. However, clinical efficacy was limited due to major challenges in delivery, longevity, and safety of synthesized siRNAs^4^.

Full chemical stabilization of the siRNA scaffold was later introduced to increase longevity of the siRNA in different tissues^5^. In addition to direct chemical modifications of siRNAs, covalent conjugations have also been explored to improve siRNA distributions in different organs^4^. Recent advancements with conjugated siRNAs have shown much more clinically relevant pharmacokinetics. Targeting liver diseases, siRNAs with a trimer of *N*-acetylgalactosamine (GalNAc) conjugate achieved efficient delivery by binding to the hepatocyte-specific Asialoglycoprotein receptor (ASGPR) showing upwards of 6-to-12-months effect in gene silencing with a single administration^6–8^. Although the platform is promising with four clinically approved conjugated siRNAs for the liver^9^, its selectivity to liver hepatocytes limits its use in other organs. Previously, the hydrophobicity of conjugates has been shown to increase accumulation and efficacy in multiple extrahepatic tissue types^10^. Similarly, increasing the molecular size by generating multivalent configurations has also been shown to increase distribution and gene silencing in the brain^11^ and lung^12^.

The first organ to be targeted by siRNA clinically was the eye^13^. Early on, non-modified or partially modified siRNAs were used resulting in limited efficacy and durability^14^. The eye is an ideal organ for siRNA treatment^9^. Its small and confined compartment allows lower drug doses to achieve a physiological improvement. Additionally, the retina is immune privileged and there are a multitude of dominant mutations that result in inherited retinal dystrophies (IRD)^15^. By far the largest number of mutations affect genes expressed in the light sensing rod photoreceptors (PR). These mutations lead to IRD that are part of a family of mutations that cause Retinitis Pigmentosa^15, 16^. Compared to rAAV mediated gene transfer, the smaller size of the siRNA offers the potential advantage to treat many of these PR diseases by intravitreal delivery. This should result in a more uniform therapeutic outcome than what can be achieved by subretinal delivery of rAAV, which results only in local transduction around the injection bleb. Since there are several retinal cell types and different tissues within the eye (Figure S1) and the structure of the conjugate and the siRNA have a profound impact on the tissue and cell distribution, we evaluated the impact of siRNA structures and conjugates on intraocular accumulation in different eye tissues and cell types, with a focus on PR enrichment. Recent work showed that conjugation of C16 to the siRNA supports efficient silencing in the retinal-pigmented epithelium (RPE) of the eye^17^. However, efficient gene silencing in retinal PRs has not yet been evaluated systematically.

To identify bioconjugates and multivalent conformations that enrich in PRs we synthesized a panel of twelve fully chemically stabilized siRNA variants, all targeting the Huntingtin (*Htt*) gene^18^, where the nature of the hydrophobic conjugate or the valency was systematically altered^10–12, 18–20^. We found that while most siRNAs variants distribute well throughout the retina, the tetravalent siRNA (hereafter referred as tetra-siRNA*^Htt^*) exhibits the best enrichment in PRs. Stability, safety, and silencing efficiency of tetra-siRNA*^Htt^*were evaluated in both, mouse, and porcine retinas. In mice, a single intravitreal injection of 15μg of the tetra-siRNA*^Htt^* is sufficient to result in a ∼75% reduction in retinal HTT protein expression for up to 6 months. In pigs, where eye size and vitreous volume are similar to humans, a single intravitreal injection of 300μg of the tetra-siRNA*^Htt^*results in a stable protein reduction of ∼80% for at least 4 months, which was the longest time point tested. Histological analyses of the retina did not reveal any long-term adverse immune reactivity in both animal models. The data suggest that therapeutic relevant gene silencing to treat PR-related retinal diseases is attainable with 1-2 intravitreal injections of chemically modified siRNA per year.

## Results

### The structure and chemical configuration of the siRNA impact distribution patterns in mouse retinas

Previous studies showed that hydrophobicity and multivalency can influence the pharmacokinetics and distribution of siRNAs^11, 12, 18, 21^. To elucidate how different bioconjugates and valances affect siRNA distribution in the retina, we synthesized twelve chemical and structural siRNA variants^10–12, 18–20^. The siRNAs were fully chemically modified, with a combination of 2’-OMe, 2’-F, vinylphosphonate, and phosphorothioates (Figure 1A). All compounds were targeting the Huntingtin (*Htt*) gene and tagged with a Cy3 fluorophore for visualization^18, 20^. We have previously established that the Cy3 dye does not principally contribute to the cellular distribution in the context of these configurations^22^. The modified chemical architectures tested included four different valences, Monovalent^12^, Divalant^11, 12^, Trivalent^12^ and Tetravalent^12^, as well as different conjugates, such as Triple-amine^20^, RA (retinoic acid)^20^, PC-RA (phosphocholine-retinoic acid)^20^, DHA (docosahexaenoic acid)^19, 20^, di-DHA (di-docosahexaenoic acid)^21^, PC-DHA (phosphocholine-docosahexaenoic acid)^20^, PC-TS (phosphocholine-α-tocopheryl succinate)^20^ and DCA (Docosanoic acid)^20, 23^ (Figure 1B). Adult mice were injected intravitreally with 1μL of each siRNA (3μg/μL) and tissues were collected 3 days post injection for section analyses. We found a predominant enrichment of siRNAs in the inner nuclear layer (INL) region, reminiscent of a Müller glia cell pattern (Figure S1), with all conjugates and valances and a relative uniform distribution of the siRNA across the entire eye (Figure 1C and 2A). Most also showed accumulation in the ganglion cell layer and in the retinal-pigmented epithelium (RPE) (Figure 1C). Accumulation in PRs, seemed to occur preferentially with the trimer, tetramer, and DCA, based on signal seen in the outer nuclear layer (ONL) and PR inner segment (IS) region. Like previous observations in the mouse brain^11^, the unconjugated monomer displayed the least siRNA retention in the retina. Nevertheless, the Cy3 signal seen, seemed to show spotty accumulation in the outer plexiform layer and where PR-ISs reside, at an interval consistent with cone PR pedicles and cone IS, respectively (Figure 1C).

**Figure 1.**
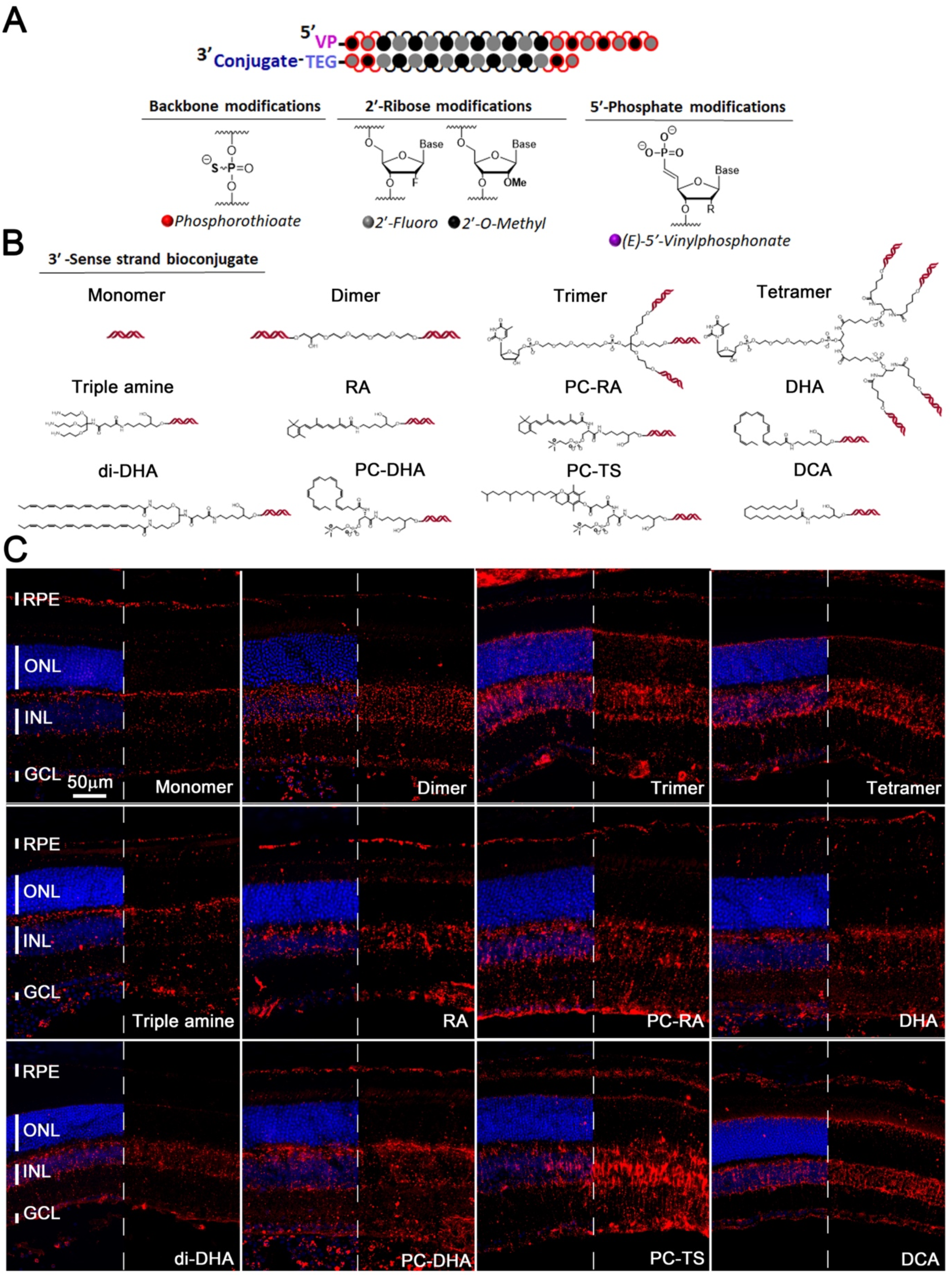
Distribution of different siRNA chemistries following intravitreal delivery. (A) Schematic of modifications on the siRNA backbone and ribose. (B) Schematics of the 12 configurations used, showing in the first row the structures of different valences and rows 2-3 the structure of the conjugates (RA: retinoic acid; PC-RA: phosphocholine-retinoic acid; DHA: docosahexaenoic acid; di-DHA: di-docosahexaenoic acid; PC-DHA: phosphocholine-docosahexaenoic acid; PC-TS: phosphocholine-α-tocopheryl succinate; DCA: Docosanoic acid). (C) Representative images of retinal cross-sections 3 days post intravitreal injection of 3μg of siRNA per eye. Valency or conjugate is indicated in each panel in the same order as shown in **b.** Blue: nuclei marked with DAPI, red: siRNAs labeled with Cy3. In each panel half of the blue signal was removed to better visualize the red signal. RPE: retinal-pigmented epithelium; ONL: outer nuclear layer; INL: inner nuclear layer; GCL: ganglion cell layer; vertical bars indicate height of different layers. Scale bar: 50μm. Histology was repeated with at least n=3 retinas.

**Figure 2.**
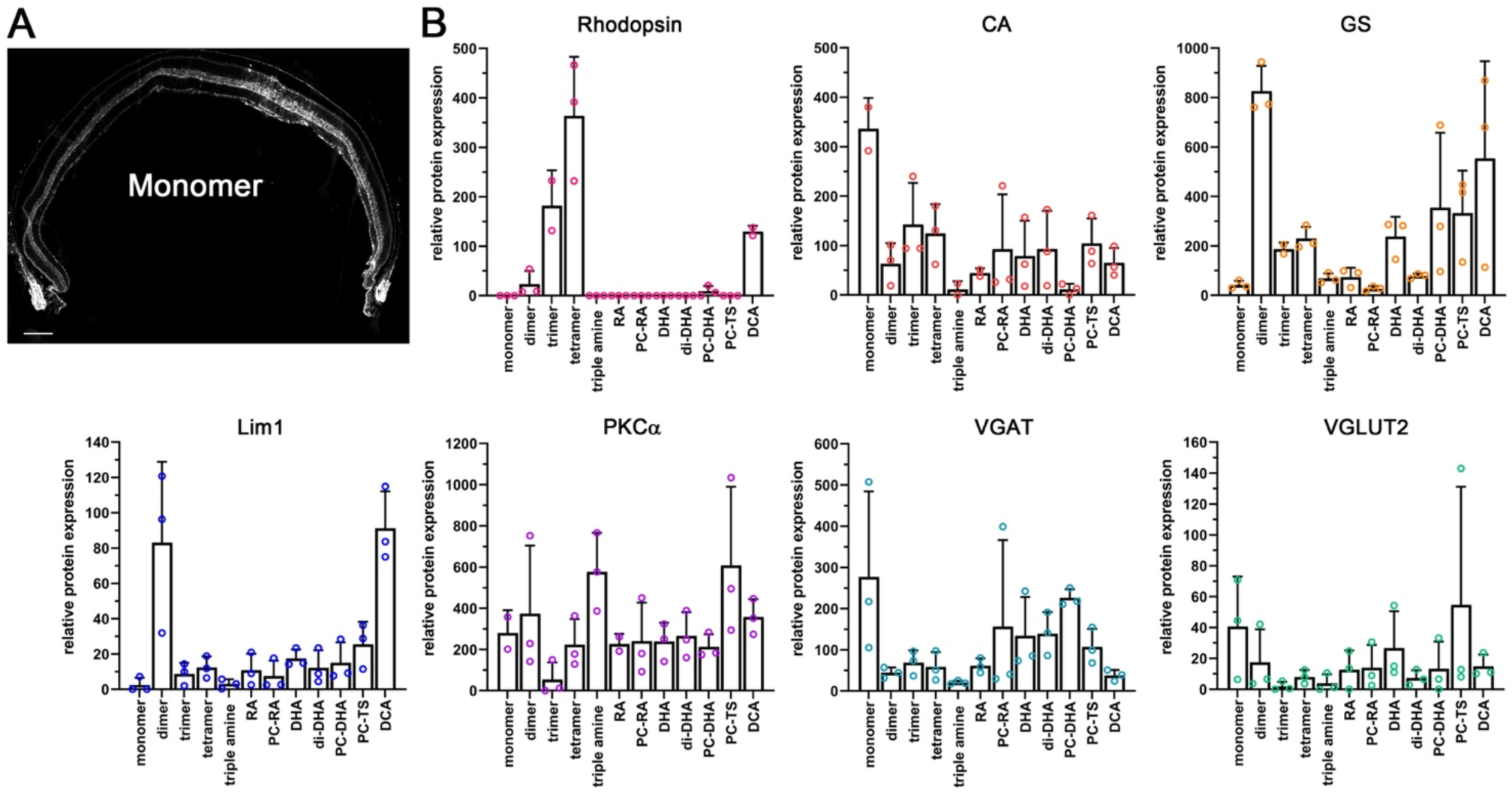
Enrichment of siRNA chemistries in different retinal cell types. (A) Cross-section showing efficient distribution of monomer siRNA labeled with Cy3 across entire retinal cross-section. Image shown in gray scale to better visualize the Cy3 signal. (B) Western blot quantifications with cell type specific antibodies from protein extracts of FACS cells that were Cy3 positive. Y-axis shows relative enrichment when compared to an un-injected retina. Antibodies: Rhodopsin (rod PRs); CA (cone arrestin: cone PRs); GS (glutamine synthetase: Müller glia); Lim1 (Lim class homeodomain transcription factor 1: mainly Horizontal cells); PKCα (protein kinase alpha: mainly rod bipolars); VGAT (vesicular GABA transporter: amacrine cells, horizontal cell); VGLUT2 (vesicular glutamate transporter 2: ganglion cells); N=3 samples; error bars: S.D..

To better identify the cell types in which the different siRNAs enrich, we repeated the injections, dissociated the retinas 3 days post-injection, and collected Cy3-labelled cells by fluorescence-activated cell sorting (FACS). Cell type specific antibodies were then used to determine by western blot, which cell types are enriched by each chemistry (Figure 2B). To capture most of the retinal cell types, we used antibodies against: rhodopsin (rod PRs); cone arrestin, CA (cone PRs); glutamine synthetase, GS (Müller glia cells); LIM class homeodomain transcription factor, Lim1^24^ (Horizontal cells); protein kinase C α, PKCα^25^ (rod bipolar cells); vesicular GABA transporter, VGAT^26^ (amacrine cells, horizontal cell); and vesicular glutamate transporter 2, VGLUT2^27^ (ganglion cells, Müller glial cells). Enrichments were seen for rods, cones, Müller glia, bipolar and amacrine/horizontal cells. As predicted by the histological analyses (Figure 1C) the multivalent chemistry and DCA were very efficient for PRs (monovalent for cones and tetravalent for rods). The strongest enrichment in Müller glia cells was seen with the divalent chemistry. Thus, similar to systemic, CNS and lung delivery, the structure and chemical composition of the siRNA has a profound impact on the intraretinal distribution profile indicating that different conjugates might be optimal for different cell types. Since the tetravalent configuration was enriched in cone PRs and displayed the strongest enrichment in rod PRs, it was selected for detailed examination of safety, distribution, and functional silencing efficiency.

### Tetrameric siRNA displays a dose dependent, safe and long-term silencing of HTT in mouse retinas

First, we evaluated the efficacy of the tetravalent siRNA at different dose levels in mice eyes. Mice were injected intravitreally with 2 μL of tetravalent siRNA to accommodate a dose range of 1 μg, 6 μg, 15 μg, 30 μg, and 60 μg per eye. Histological analyses at 2-months post-injections revealed a uniform distribution of the Cy3-labelled siRNA, with a signal intensity proportional to the injected dose (Figure 3A). Gene silencing by the previously optimized huntingtin (*Htt*) sequence^18^ was evaluated by western blot. We found a dose dependent reduction in HTT protein (Figure 3B), with a silencing efficiency of ∼70% at around 15μg of tetra-siRNA*^Htt^*. The non-targeting control (NTC), which consists of a scrambled sequence of the same chemical configuraiton^18^, showed no reduction in HTT protein at 15μg, indicating the specificity of the observed silencing activity (Figure 3B).

**Figure 3.**
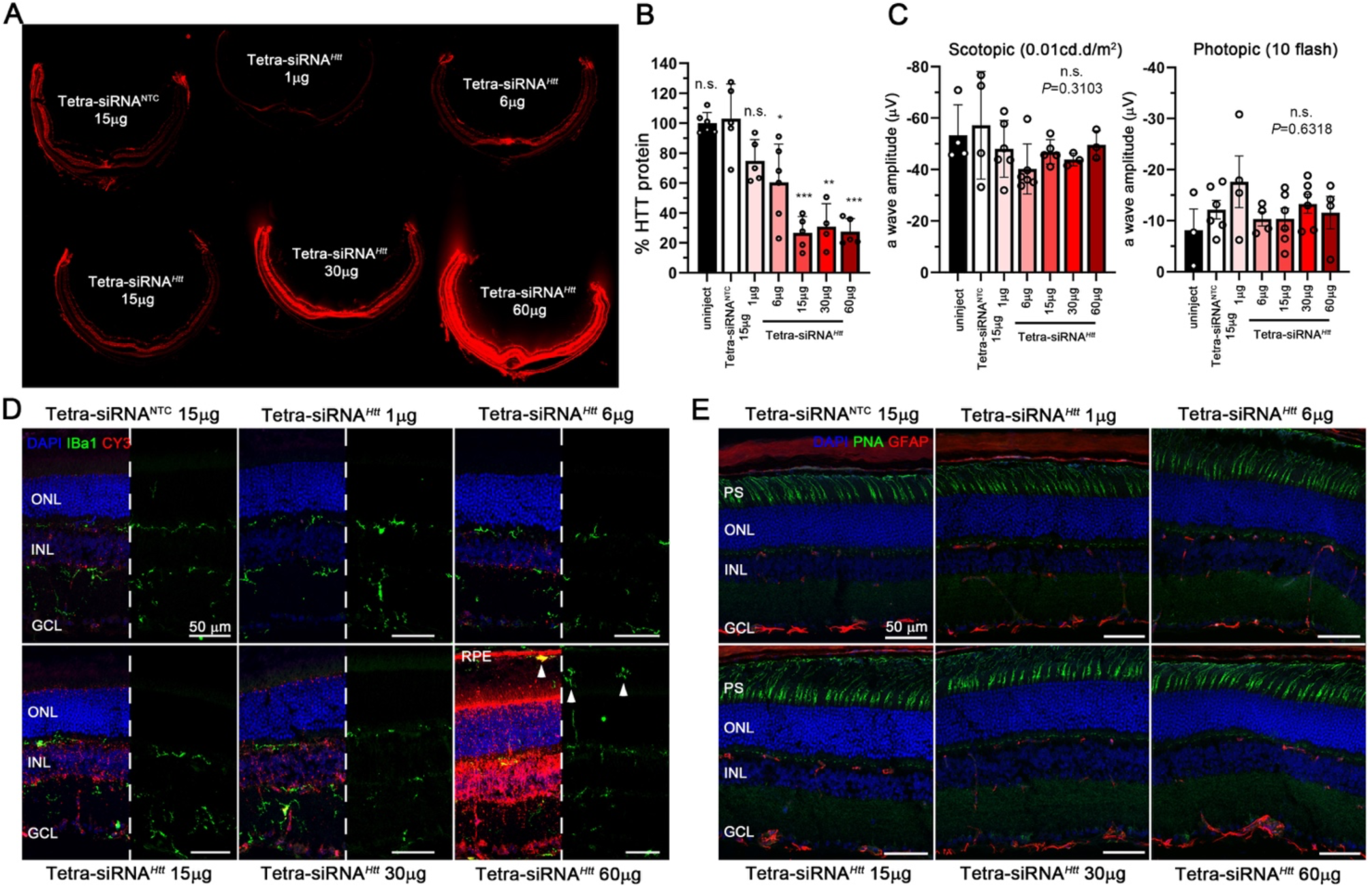
Dose escalation of tetravalent siRNA in mouse. (A) Retinal cross-sections of mice 2 months post-injection of 2μL siRNA (doses indicated in section). Seen is the siRNA distribution through the fluorescence of the attached Cy3 fluorophore. (B) Dose response at 2 months post-injection using same doses and injection volume as shown in panel (A). Shown is percentage of remaining HTT protein compared to the NTC. (C) Scotopic (first graph) and photopic (second graph) a-wave amplitudes from ERG recordings at 2 months post injection (N=6 for (A-C), error bars: S.D.; significance is compared to NTC; n.s.: not significant; *: p<0.05; **: p<0.01; ***: p<0.001). (D, E) Retinal cross-sections shown in panel (A) stained for Iba1 (D: green signal) and GFAP (E: red signal) to visualize migrating microglia and activation of gliosis, respectively. Except for the highest dose (60μg) there are no microglia (white arrowheads) seen in the outer nuclear layer or where PR segments reside. Blue: nuclei marked with DAPI; green: Iba1 in panel (D) and peanut agglutinin lectin (PNA) marking cone PR segments in panel (E); red: siRNA in panel (D) marked through Cy3-label and GFAP in panel **e**. In each panel of **d** half of the blue was removed to better visualize the red and green signal. Histology in (A, D, E) was repeated with at least n=3 retinas. PS: photoreceptor segments; ONL: outer nuclear layer; INL: inner nuclear layer; GCL: ganglion cell layer; RPE: retinal-pigmented epithelium; white arrowheads mark microglia in the subretinal space; vertical bars indicate height of different layers. Scale bars: 50μm.

To evaluate safety, we performed electroretinogram (ERG) recordings under scotopic (rod PR response) and photopic (cone PR response) conditions, which revealed no significant differences among the different dose groups (Figure3C). Similarly, immunohistochemical analyses on retinal cross-sections detected no migration of microglia marked by ionized calcium binding adaptor molecule 1 (Iba1) into the ONL and/or inner outer segment region, except for occasional cells at the highest dose (60μg, arrowheads) (Figure 3D). Consistent with this, there was no reactive gliosis in the retina as assessed by the normal expression of glial fibrillary acidic protein (GFAP) (Figure 3E). Both, Iba1 and GFAP are markers to assess early retinal immune responses and early signs of neurodegeneration^28^. Together, the data indicate an efficient and safe gene silencing by the tetra-siRNA*^Htt^*in the mouse retina.

To examine the longevity of tetra-siRNA*^Htt^*, mice were intravitreally administered a medium dose (15μg/eye) of tetra-siRNA*^Htt^* and silencing efficiency was evaluated 3-, 6- and 9-months post-injection. The tetra-siRNA*^NTC^* was used as a control. At three-month post injection, the tetra-siRNA*^Htt^*was easily visualized by fluorescent microscopy on retinal cross-sections. Uniform distribution throughout retina over time was apparent by fluorescent funduscopy (Figure 4A). Section analyses for HTT protein expression showed a strong reduction of HTT protein throughout the retina including PR-ISs, showing that the tetravalent siRNA does indeed target PRs efficiently (Figure 4B). Similarly to the dose escalation study, where there was no inflammation at 2-months post-injection with a dose of 15μg, there was no change in GFAP (Figure 4C) and Iba1 (Figure 4D) expression at all 3 time points. Western blot quantification of HTT protein showed a robust reduction of ∼75% and ∼70% of HTT protein at 3- and 6-months post-injection, respectively. However, by 9 months post-injection there was no significant differences between the two groups (Figure 4E). In summary, the mouse data indicate that PR-expressed genes can be efficiently downregulated with the tetravalent siRNA in a safe manner over a prolonged time-period.

**Figure 4.**
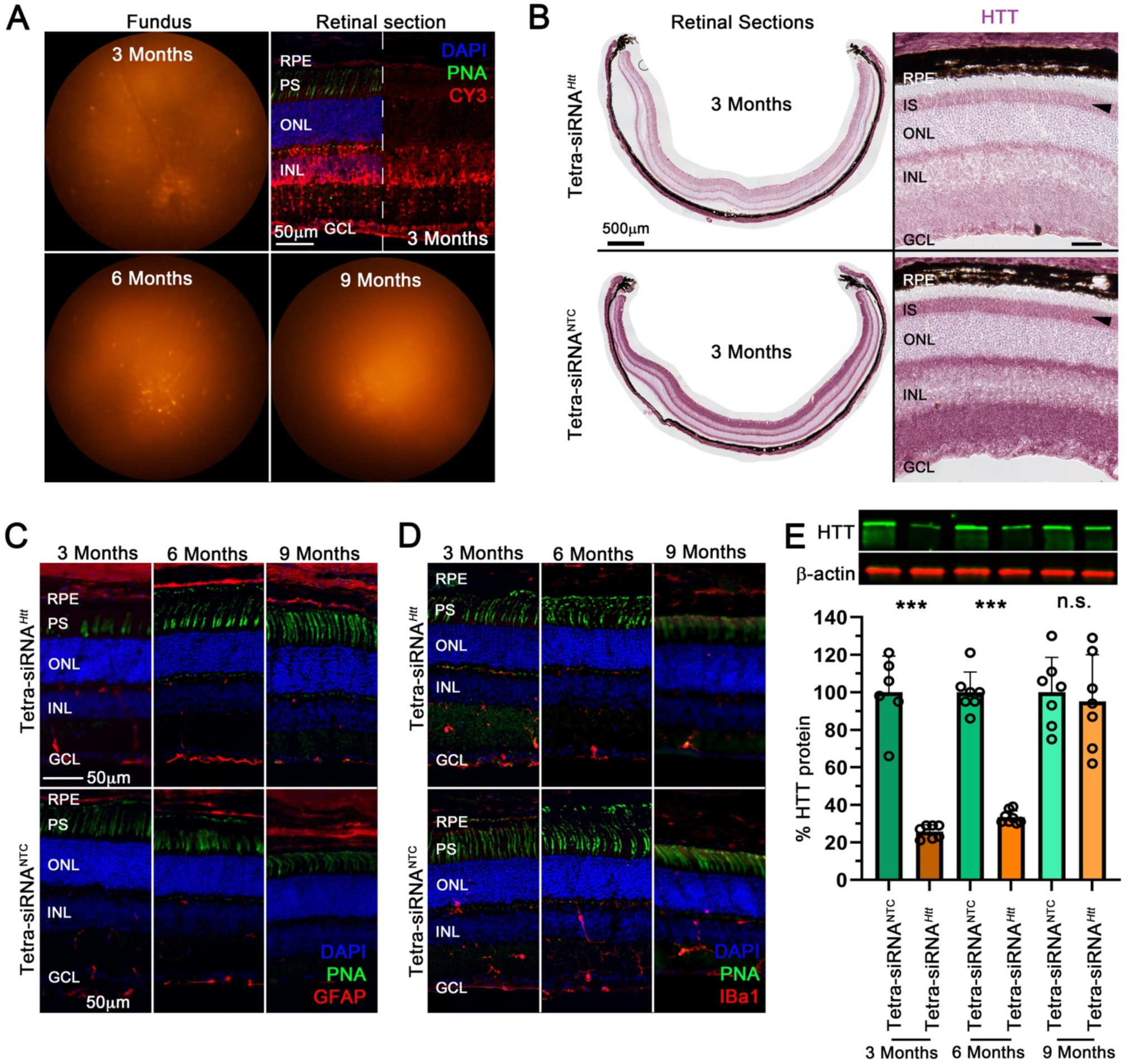
Long-term silencing of HTT with tetravalent siRNA. (A) Fluorescent fundus images and retinal cross-section of mice injected with 15μg of the tetra-siRNA*^Htt^* and imaged at time points indicated to show the Cy3 signal of the siRNA. (B) Retinal cross-sections at 3 months post-injection stained for expression of the HTT protein (purple signal). Shown is an entire cross-section (left panels) and a magnified view (right panels). Reduction of HTT protein in PR inner segments (IS) is indicated by black arrowheads. (C, D) Retinal cross-sections stained for GFAP (red signal) and Iba1 (red signal) at 3-, 6-, and 9-months post intravitreal injection with 15μg siRNAs to visualize activation of gliosis and migration of microglia, respectively. No gliosis and migration of microglia is seen over time at the dose used. (E) Quantification of remaining retinal HTT protein over time after one intravitreal injection of 15μg siRNAs. Top shows example of protein bands on western blot and bottom shows quantification. Shown is percentage of remaining HTT protein compared to the NTC for each time point (N=6-7 retinas per time point and siRNA type; error bars: S.D.; n.s.: not significant; ***: p<0.001). (A, C, D) Blue: nuclei marked with DAPI; green: cone segments marked with peanut agglutinin lectin (PNA); red: siRNAs in panel (A) marked by Cy3-label and GFAP and Iba1 in panels (C, D). In panel (A) half of the blue was removed to better visualize the red signal. RPE: retinal-pigmented epithelium; PS: photoreceptor segments; IS: inner segment; ONL: outer nuclear layer; INL: inner nuclear layer; GCL: ganglion cell layer. Scale bars: 50μm except where indicated otherwise in panel (B). Histology in (A-D) was repeated with at least n=3 retinas.

### Tetra-siRNA^HTT^ delivery in porcine retina

In mouse the lens is ∼20% larger than the vitreous, which is in contrast to the human anatomy where the lens volume is ∼16 times smaller than the vitreous volume. Additionally, the human vitreous is ∼500 times larger than the mouse vitreous. These differences in relative and absolute sizes may alter how siRNAs behave in large eyes. To test the suitability of the tetravalent siRNA for clinical use we injected the tetra-siRNA*^Htt^* intravitreally into the dorsal-temporal sclera (Figure 5A) of porcine eyes^29^. An initial dose escalation (100μg, 500μg, 1000μg, 1500μg) was used to determine distribution, toxicity and silencing efficiency over a period of 2 weeks (Figure 5B-D). Upon enucleation the cone-dominant central visual streak was processed for histology (Figure 5A), while the remaining peripheral quadrants (Figure 5A) were used to quantify the silencing of HTT protein (Figure 5D). Distribution of the Cy3-labelled tetra-siRNA*^Htt^*appeared dose and injection site dependent (Figure 5B), which was confirmed by western blot analyses of the 4 quadrants within one eye (Figure 5D). The average silencing efficiency increased only by ∼10% between 500μg (∼60% silencing) and 1500μg (∼70% silencing), suggesting a possible saturation at around 500μg/eye. Histological analyses revealed an increase in GFAP expression and Iba1 positive microglia starting at 1000 μg/eye, indicating that the higher doses cause retinal toxicity (Figure 5C). At 500 μg/eye only a moderate upregulation of GFAP in Müller glial was seen and no migration of microglia into the ONL, suggesting that doses below 500μg/eye are safe for large eyes.

**Figure 5.**
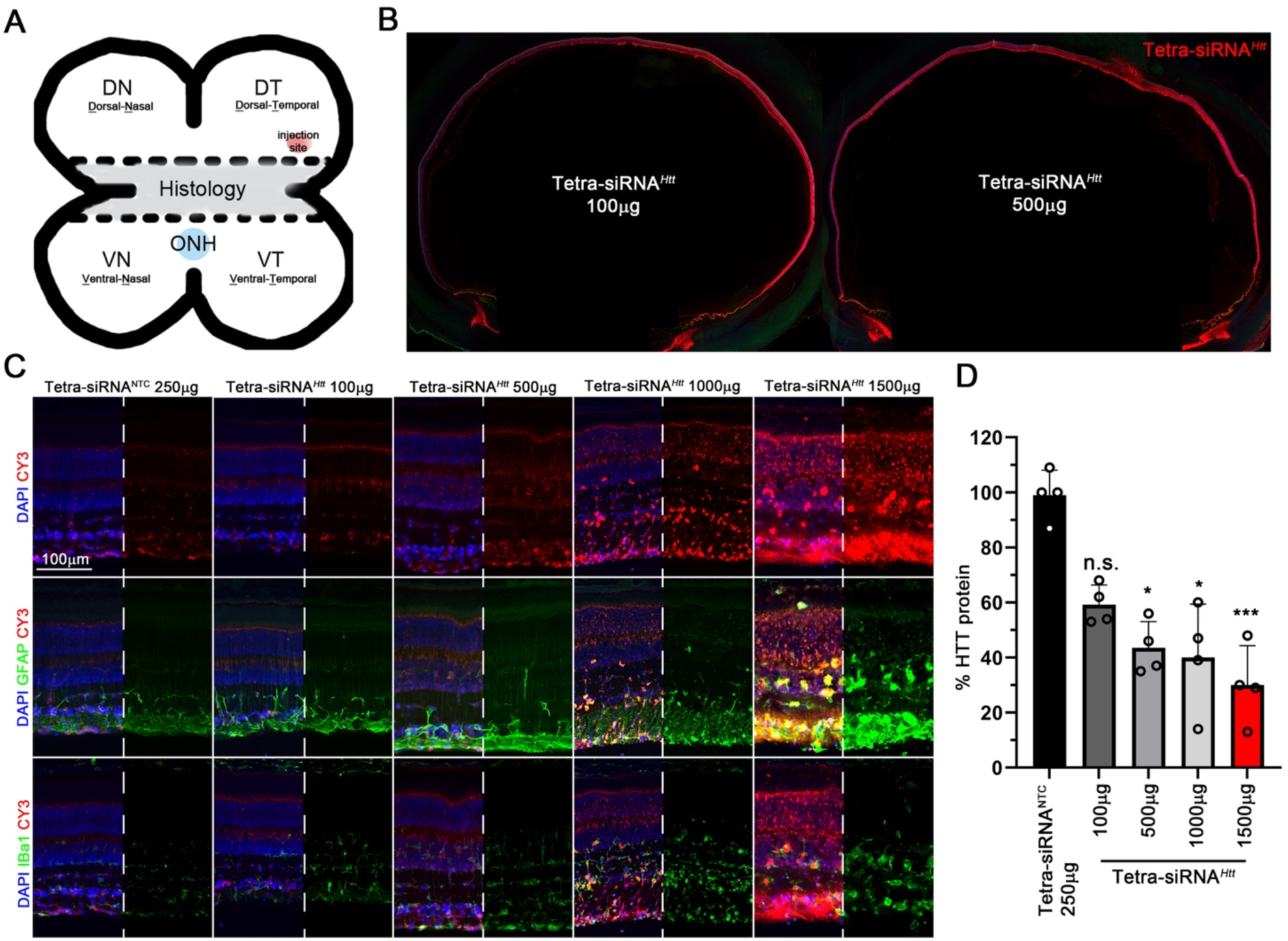
Dose escalation of tetravalent siRNA in porcine eyes. (A) Schematic of how porcine eyes were processed and where eyes were injected. The central part was used for histological analyses and the dorsal and ventral parts of the eye were used for quantitative analyses. ONH: Optic nerve head. (B) Distribution of Cy3-labeled siRNA across the eye 14 days post intravitreal injection at doses indicated. (C) Retinal cross-section of eyes 14 days post intravitreal injection with siRNAs. Doses are indicated on top of each column. Sections were stained for GFAP (middle row) or Iba1 (last row) to visualize activation of gliosis and migration of microglia, respectively. Blue: nuclei marked with DAPI; green: GFAP or Iba1 as indicated; red: siRNAs marked by Cy3-label. In each panel half of the blue signal and for rows 2 and 3 also the red signal, were removed to better visualize the green signal in rows 2 and 3 and the red signal in row 1. ONL: outer nuclear layer; INL: inner nuclear layer; GCL: ganglion cell layer. Scale bar: 100μm. (D) Percentage quantification of remaining retinal HTT protein when compared to NTC at 14 days post intravitreal injection at doses indicated in bar graph. Each quantification represents the average of the 4 quadrants of one eye (error bars=S.D.; significance is compared to NTC; n.s.: not significant; *: p<0.05; ***: p<0.001).

To assess silencing efficiency and toxicity over time we injected a dose deemed safe from the dose escalation study (300μg/eye) and analyzed the tissue at 1- and 4-months post-injection. Histological analyses showed a clear reduction of HTT protein that was maintained over a period of 4 months (Figure 6A). In particular, the reduction in PR-ISs was clearly visible (Fig. 6a: arrows). The histological data was corroborated by western blot analyses, which showed a ∼80% reduction in HTT protein over the duration of the 4-months (Figure 6B). To understand the pharmacokinetics of the siRNA we measured its distribution within the different tissues of the eye by performing a peptide nucleic acid (PNA) hybridization assay^30^ (Figure 6C). The PNA assay evaluates the presence of the guide strand, which is not dependent on the presence of the fluorescent label. The highest concentration of siRNA was found in the retina followed by the RPE. The vitreous, cornea and lens had detectable amounts of the oligonucleotide, which was at least fifty-fold lower (Figure 6C). Interestingly, in the RPE the concentration of the siRNA stayed stable between 1-4 months post - injection, while it declined in all other tissues. Finally, analyses of Iba1 and GFAP expression revealed few microglia migrating into subretinal space close to the injection site at 1-month post-injection, however, this transient migration was not observed at 4-month post-injected (Figure S2) Similarly, there was mild gliosis near the injection site at 1-month but not at 4-months post-injections, as seen by changes in GFAP expression (Figure S2). In summary, the data suggest that safe, efficient, and long-lasting silencing of PR-expressed genes is feasible as a therapy in humans with the tetravalent siRNAs.

**Figure 6.**
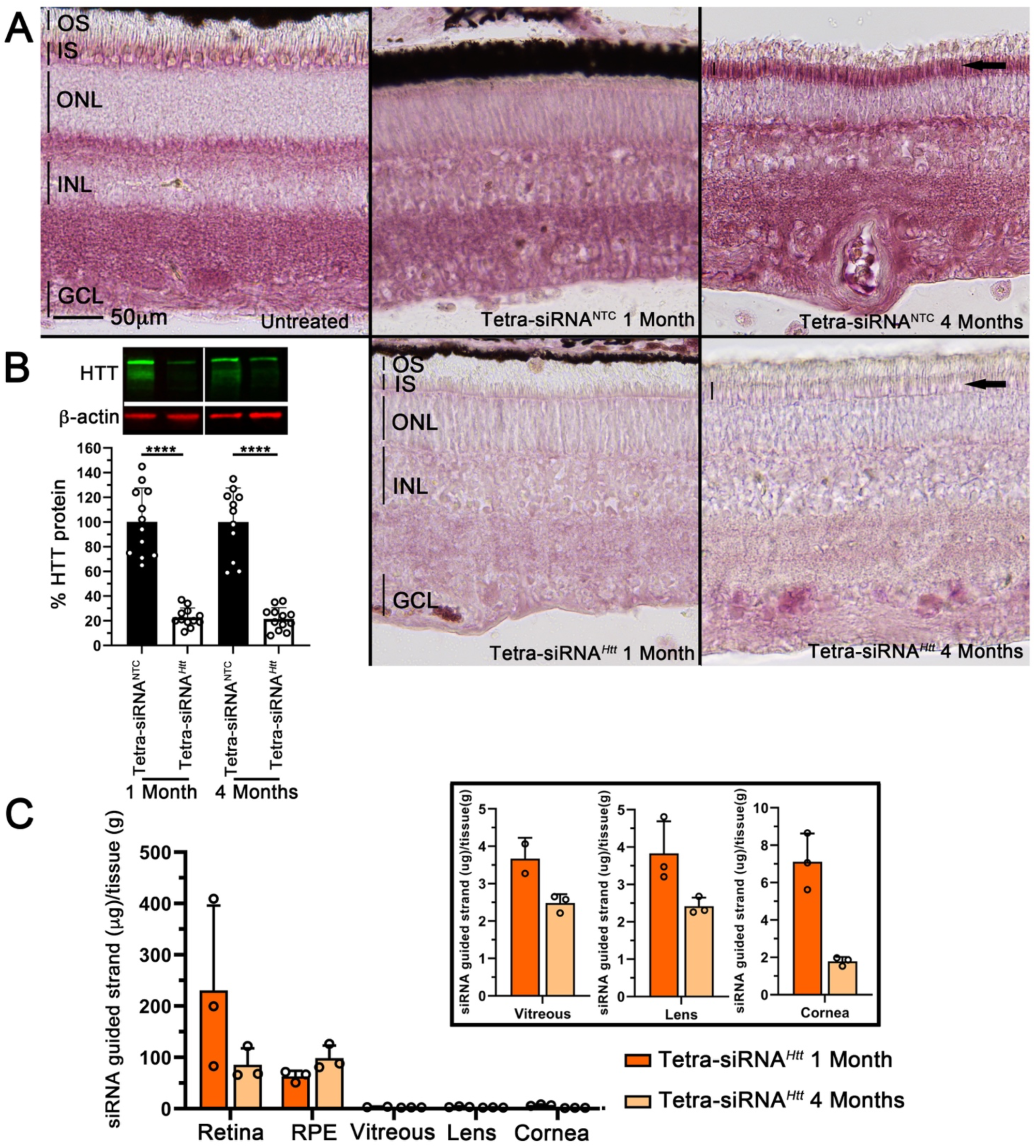
Long-term HTT silencing with tetravalent siRNA in porcine retina. (A) Retinal cross-section stained for expression of the HTT protein (purple signal). Top row shows un-injected or non-targeting control (NTC) injected eyes at 1- and 4-months post-injection with 300μg of siRNA. Bottom row shows tetra-siRNA*^Htt^*injected eyes with same dose. Silencing in PR inner segments (IS, arrows and vertical bars marking height of IS in last column) is still clearly visible at 4-months post-injection. Images are representative images from 3 pigs per time point and siRNA. OS: outer segments; IS: inner segments; ONL: outer nuclear layer; INL: inner nuclear layer; GCL: ganglion cell layer. Scale bar: 50μm. (B) Percentage quantification of remaining retinal HTT protein at 1- and 4-months post-injection when compared to the NTC. Top image shows examples of bands on western blot gel. Quantification represents the average of the 4 quadrants of 3 eyes (N=12; error bars: S.D.; ****: p<0.0001). (C) Pharmacokinetics of remaining siRNA in different eye tissues. PNA (peptide nucleic acid) hybridization assay was performed with one quadrant per eye (N=3, error bars: S.D.). Analyses of the retina, RPE (retinal-pigmented epithelium), vitreous, lens and cornea show that most of the siRNA is taken up by the retina and RPE and cleared from the vitreous. Additionally, very little migrates to the lens and cornea. Eye-tissues from NTC injected eyes did not show any signal and were therefore omitted from the figure. Boxed area shows vitreous, lens and cornea graphs with adjusted Y-scales.

## Discussion

Here, we examined 12 different modified siRNAs and their retinal cell distribution. We show that while all of them distribute well across the retina, the nature of the siRNA structure and the conjugate has a significant impact on the preference of the cellular distribution. One interesting finding was that none of the lipid conjugates showed any preferential enrichment in PRs. In contrast, tertravalent siRNA did show PR enrichment and was therefore evaluated further for silencing efficiency, safety and longevity in mouse and porcine retinas. The histological analyses in mouse and porcine showed that the tetravalent siRNA is quite effective at causing HTT silencing in PRs. In mouse the silencing appears to last between 6-9 months. While we did not determine the silencing duration in porcine retinas past 4 months, it is safe to assume from the data collected that a similar if not longer time window applies to the porcine retina. This is therapeutically relevant since depending on the disease that is being treated 1-2 intravitreal injections per year may be sufficient to have a significant therapeutic effect. Many IRD-related genes are associated with dominant mutations in PR-expressed genes, meaning that silencing the mutated copy of the gene may be sufficient to significantly delay disease progression. siRNAs represent an ideal approach for this type of mutations. The most significant advantage of the siRNA approach over the rAAV approach in targeting PRs is the intravitreal delivery route. While being less risky and invasive it also results in widespread targeting of all PRs. This is likely to result in a better therapeutic outcome over time, when compared to the local PR transduction that is achieved with rAAVs. Additionally, once a chemistry is selected and tested for safety, the only changes needed to treat a different dominant IRD are changes in a few nucleotides of the siRNA sequence. This can significantly increase the speed and decrease the cost at which therapies can be developed, making eye saving injections available to the many who still suffer from these debilitating diseases.

Many non-IRDs such as diabetic retinopathy (DR), diabetic macular edema (DME) and the exudative form of age-related macular degeneration (AMD), could also benefit from long lasting siRNA therapies. All these diseases are caused by abnormal neovascularization. Currently these diseases are treated with anti-vascular endothelial growth factor (VEGF) inhibitors requiring intravitreal injections at a frequency of every 4-8 weeks^31^. To date, there are two clinical trials using siRNA to treat these diseases^32–35^, however, these trials were stopped due to a failure to reach the primary endpoint. Our divalent chemistry enriched well in Müller glia cells and to some extend in the RPE. Both cell types are believed to be important target cell in reducing retinal VEGF levels in these diseases^36^. Similarly, a new FDA approved drug that delays disease progression in the dry form of AMD, targets the complement system in RPE cells^37, 38^ with an injection frequency of every 4-8 weeks. The frequency of injections for these age-related illnesses is a burden to elderly patients. Any therapeutic approach that reduces the intra ocular injection frequency constitutes a significant improvement in the quality of life for these patients. In summary, by taking a cell type specific approach we have identified chemistries for efficient transduction of Müller glia cells and PRs. These chemistries represent new opportunities as to how age-related as well as IRDs can be treated in the future. Additionally, they increase speed and reduce costs for the development of these sight saving treatments.

## Materials & Methods

### Animals

All procedures involving animals were in compliance with the Association for Research in Vision and Ophthalmology (ARVO) Statement for the Use of Animals in Ophthalmic and Vision Research and were approved by the Institutional Animal Care and Use Committees (IACUC) of the University of Massachusetts Medical School. Mice used in the study were between the ages of 2-3 months at the time of injection and were C57BL/6J mice purchased from the Jackson Laboratory. Pigs used in the study were between the ages of 1-2 months at the time of injection and were Yorkshire pigs purchased from Earl Parsons and Sons Inc.

### Oligonucleotide synthesis

Oligonucleotides were synthesized by phosphoramidite solid-phase synthesis on a MerMade12 (Biosearch Technologies, Novato, CA) or on AKTA Oligoplilot 10 or 100 (Cytiva, Marlborough, MA) using 2ʹ-F or 2ʹ-O-Me modified phosphoramidites with standard protecting groups. 5’-(E)-Vinyl tetra phosphonate (pivaloyloxymethyl) 2’-O-methyl-uridine 3’-CE phosphoramidite (VP) was purchased from Hongene Biotech, USA, Cy3 phosphoramidite (Quasar 570 CE) was used to label the oligonucleotides purchased from GenePharma, Shanghai, China. Trivalent and Tetravalent oligonucleotides were prepared using commercial trebler and doubler phosphoramidites respectively as branching points, (Glen Research, Sterling, VA) and Tetraethyloxy-Glycol phosphoramidite (TEG) (ChemGenes, Wilmington, MA) as spacer from the solid support. Phosphoramidites were prepared at 0.1 M in anhydrous acetonitrile (ACN), except for 2’-O-methyl-uridine phosphoramidite dissolved in anhydrous ACN containing 15% dimethylformamide. 5-(Benzylthio)-1H-tetrazole (BTT) was used as the activator at 0.25 M and the coupling time for all phosphoramidites was 4 min, except for trebler, doubler, and TEG phosphoramidites, where coupling time used was 8 min. Detritylations were performed using 3% trichloroacetic acid in dichloromethane (MerMade12) or in Toluene (AKTA). Capping reagents used were CAP A (20% n-methylimidazole in ACN) and CAP B (20% acetic anhydride and 30% 2,6-lutidine in ACN). Phosphite oxidation to convert to phosphate or phosphorothioate was performed with 0.05 M iodine in pyridine-H2O (9:1, v/v) or 0.1 M solution of 3-[(dimethylaminomethylene)amino]-3H-1,2,4-dithiazole-5-thione (DDTT) in pyridine (ChemGenes) for 3 min. Reagents for detritylation, iodine oxidation and capping were purchased from AIC. Unconjugated oligonucleotides were synthesized on 500Å (or 1000Å for Trivalent and Tetravalent oligonucleotides) long-chain alkyl amine (LCAA) controlled pore glass (CPG) functionalized with Unylinker terminus (ChemGenes). Divalent oligonucleotides (DIO) were synthesized on modified solid support prepared in house^11^. DCA, DHA, RA, PC-DHA, PC-RA, PC-TS, diDHA, and triple amine, bioconjugated oligonucleotides were synthesized on modified solid supports prepared in house^19–21^.

### Deprotection and purification of oligonucleotides for in-vivo experiments

Cy3 labeled bioconjugated oligonucleotides were cleaved and deprotected in 28-30% ammonium hydroxide-40% aq. methylamine (1:1, v/v) (AMA) for 2 hours at room temperature. Cy3 labeled multivalency oligonucleotides (mono, divalent, trivalent and tetravalent) were cleaved and deprotected by AMA treatment for 2 hours at 45°C. The VP containing oligonucleotides were cleaved and deprotected as described previously^39^. Briefly, CPG containing VP-oligonucleotides was treated with a solution of 3% diethylamine in 28-30% ammonium hydroxide at 35⁰C for 20 hours. All solutions containing cleaved oligonucleotides were filtered to remove the CPG and dried under vacuum. The resulting pellets were re-suspended in 5% ACN in water. Purifications were performed on an Agilent 1290 Infinity II HPLC system. VP oligonucleotides were purified using a custom 20×150mm column packed with Source 15Q anion exchange resin (Cytiva, Marlborough, MA); running conditions: eluent A, 10 mM sodium acetate in 20% ACN in water; eluent B, 1 M sodium perchlorate in 20% ACN in water; linear gradient, 10 to 35% B in 40 min at 50°C. Cy3 labeled multivalency and bioconjugated oligonucleotides were purified using a 21.2×150mm PRP-C18 column (Hamilton Co, Reno, NV); running conditions: eluent A, 50 mM sodium acetate in 5% ACN in water; eluent B, 100% ACN; linear gradient varied due to hydrophobicity differences of the bioconjugates in the range 10 to 25-70% B in 40 min at 60°C. Flow was 30mL/min in both methods and peaks were monitored at 260 nm for non-labeled oligonucleotides and 550 nm for Cy3 labeled oligonucleotides. Fractions were analyzed by liquid chromatography mass spectrometry (LC–MS), pure fractions were combined and dried under vacuum. Pure oligonucleotides were re-suspended in 5% ACN and desalted by size exclusion on 25×250 mm custom columns packed with Sephadex G-25 media (Cytiva, Marlborough, MA), separate columns were used for non-labeled and Cy3 labeled oligonucleotides. Desalted oligonucleotides were finally lyophilized.

### PNA Hybridization Assay

siRNA accumulation was quantified as described previously using PNA hybridization assay^30^. Harvested pig eye tissue was weighed and lysed in 300 μL homogenizing buffer (Affymetrix, #10642) using a QIAGEN TissueLyser II (Qiagen) except for cornea which was lysed in 1 mL using Bio-Gen PRO200 Homogenizer (Proscientific 01-01200).

### Cell Collection Using Flow Cytometry

Retinas from 2 injected mice for each of the different Cy3-labelled siRNAs were pooled and dissociated into single cells using papain for dissociation according to the manufacture’s instructions (Worthington; Cat# 9035-81-1). Each conjugate was done in triplicates with a total of 6 mice per configuration. Cy3*+* cells were collected by fluorescence-activated cell sorting (FACS) for western analysis to measure which modification enrich for which cell type specific protein.

### Intravitreal Administration of siRNAs in Mice and Swine

Intravitreal injections in mouse were performed as previously described^40^. The same glass needles (Clunbury Scientific LLC; Cat no. B100-58-50) were used in combination with the FemtoJet (Eppendorf) with a constant pressure and time to deliver ∼1-2ul of siRNA into the vitreous. Depending on the experiment, various amounts of siRNA were used as noted in the text. Intravitreal injections in pigs were done under anesthesia, which was performed and supervised by animal medicine of the University of Massachusetts Chan Medical School. After pigs were fully anesthetized, proparacaine and iodine were used to num and disinfect the eye, respectively. A sterile insulin syringe was used to deliver 100μL of siRNA into each eye. The injection sites were always ∼2-3mm away from the limbus on the temporal side of the eye. The injection was performed like an intravitreal injection for anti-VEGF drugs. After injection, sterile saline was used to rinse the eye. Injected amounts are described in the text.

### Funduscopy & Electroretinography

Funduscopy was performed as previously described^41^. In addition, to regular brightfield visualization the TRITC filter was used to visualize the retention of the Cy3-labelled siRNAs in the retinas. Images were acquired with the Micron IV System from Phoenix Technology Group. Electroretinograms (ERGs) were performed as previously described^41^ in mice that were injected with the doses indicated in figure 2c, at 2 months post-injection. ERGs were performed with the Celeris System (Diagnosys LLC) and their preset programs for scotopic and photopic recordings. Data shown represent the average of 6-7 mice and were recorded with the following parameters. Scotopic recordings were performed at 1 cd.s/m^2^. Photopic ERG recordings used a background intensity of 9cd.s/m^2^ and a flash intensity of 10 cd.s/m^2^.

### Histology

Mouse retinal dissection, histological preparation and immunohistochemistry staining procedures were performed as previously described^41^. The following antibodies were used: HTT (1:300; Cell Signaling, Cat# 5656), fluorescein peanut agglutinin lectin (PNA) (1:1,000; Vector Laboratories, Cat# FL1071), rabbit anti-Iba1 (1:300; Wako, Cat#019-19741), mouse anti GFAP (1:1,000; Millipore, Cat#MAB5628), rhodamine phalloidin (1:1,000; Life Technologies, Cat# R415). Nuclei were counterstained with 4’, 6-diamidino-2-phenylindole (DAPI) (Sigma-Aldrich, Cat# 9542). siRNAs were visualized by their Cy3-signal without signal enhancement. Porcine histological procedures used the same method as for mice with the following modification. From each eye, a ∼10mm wide horizontal band in the nasal-temporal direction containing the visual streak was cut from the center of the eye (Fig. 4a) and fix in 4% paraformaldehyde for 24hr. Thereafter the tissue was embedded in optimal cutting medium (OCT) and processed as described for mouse tissue.

### Quantitative Western Blot Analyses

Collected tissue were flash frozen prior to conduct the western blot analyses. Sample preparation and western blotting were performed as previously described^41^. Sample sizes are as indicated in figure legends. For pig eyes, the remaining of the retina were split into four quadrants based on the orientation. In each quadrant, a further incision was made to split the tissue into half, with one half using for western analyses. Antibodies used for protein level detections are: mouse anti b-actin (1:1,000; Cell signaling, Cat#3700), rabbit anti HTT (1:1,000; Cell signaling, Cat#5656), rabbit anti PKCa (1:1,000; Cell signaling, Cat#59754), mouse anti Rhodopsin (1:1,000; Invitrogen, Cat#MA1-722), rabbit anti Cone Arrestin (1:1,000; Millipore, Cat#AB15282), mouse anti GS (1:1,000; Millipore, Cat#MAB302), rabbit anti Lim1 (1:1,000; Invitrogen, Cat#PA5-116485), rabbit anti VGAT (1:1,000; Invitrogen, Cat#PA5-27569), VGLUT2 (1:1,000; Cell signaling, Cat#71555).

## Data Availability

All data and experimental parameters related to this paper are available in the main text and the materials and methods.

## Acknowledgments

This work was supported in part by a NEI grant (R01EY032461) and a pilot award from the UMass Center for Clinical and Translational Science with funding from NCATS/NIH (Award # UL1TR001453) to CP and an R35 GM131839 and S10 OD020012 to AK.

## Author Contributions

S.-Y.C., J.C., A.B., J.F.A., D.E., N.H., M.H., S.J., D.G., J.C. and C.P. performed experiments and interpreted data. S.-Y.C., J.F.A., A.K and C.P. conceived experiments and wrote the manuscript.

---

**Figure S1.**
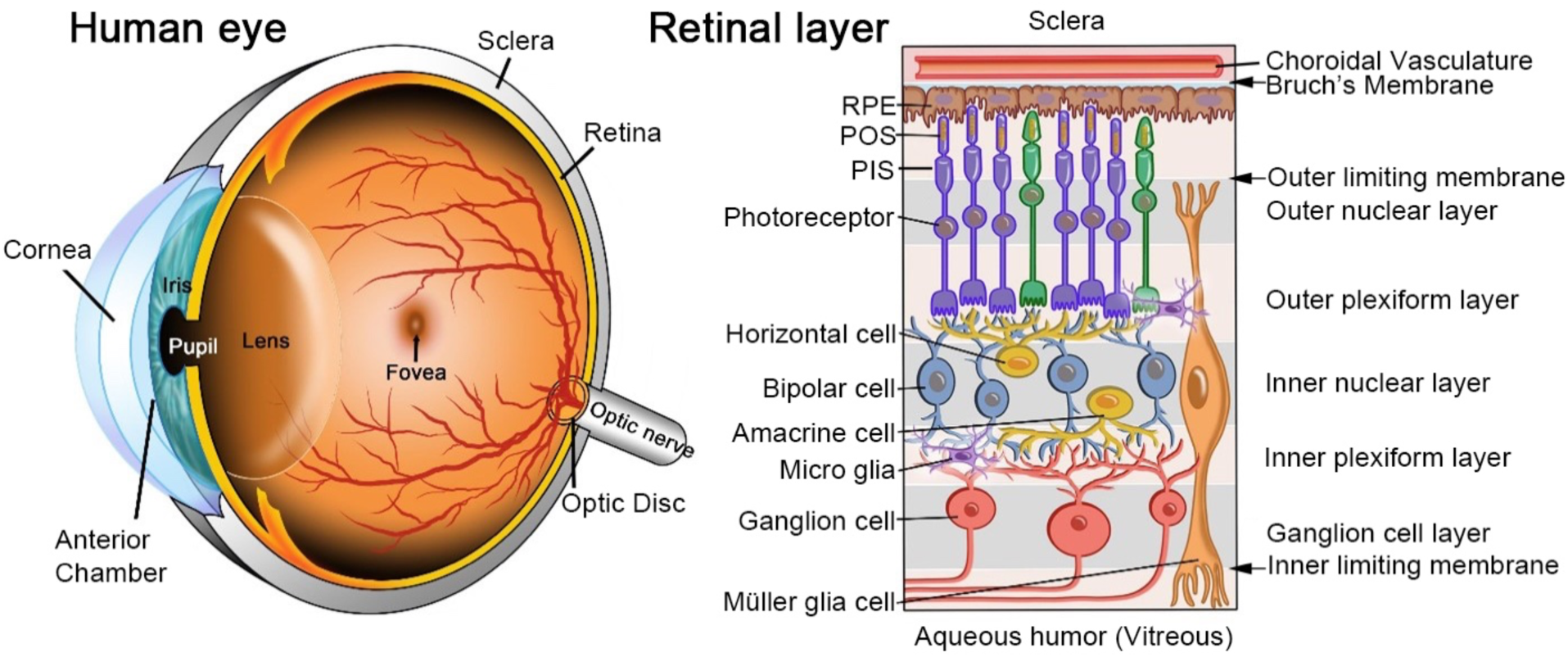
Schematic of eye and retina. Left panel: Schematic of a human eye showing the cornea, the anterior chamber, which is the space between the cornea and the iris, the pupil, the lens, the retina with the fovea, which is the area of high acuity vision in humans, the optic disc and the optic nerve, and the tissue holding the eye together, the sclera. Right panel: Schematic of a retinal cross-section showing the different cell layers and cell types. Width of individual layers are not to actual proportions. The Müller glia cells, span the retina, have the nucleus in the inner nuclear layer and form with their end-feet the inner and outer limiting membranes. Ganglion cells are in the ganglion cell layer and project their axons through the optic nerve to the brain. All interneurons are in the inner nuclear layer, including amacrine cells, horizontal cells, and bipolar cells. Amacrine and bipolar cells connect to ganglion cells in the inner plexiform layer while horizontal cells and bipolar cells connect to photoreceptors in the outer plexiform layer. Photoreceptors have their cell bodies in the outer nuclear layer. The photoreceptor inner segment (PIS) is on the outer side of the outer limiting membrane and contains all the major cytoplasmic components of the cell. The photoreceptor outer segment (POS) is connected to the PIS by the connecting cilium. The POS is the region where light photons are absorbed. POSs are surrounded by apical microvilli processes of the retinal pigmented epithelium (RPE). The RPE digests every day ∼10% of each POS. RPE cells are attached to a basal membrane referred to as the Bruch’s membrane. On the other side of the Bruch’s membrane is the choroidal vasculature and the sclera. The siRNA that is injected intravitreally migrates across the inner limiting membrane and the retinal cell layers to the photoreceptors and the RPE cells. Very little remains in the vitreous or migrates to the lens and cornea.

**Figure S2.**
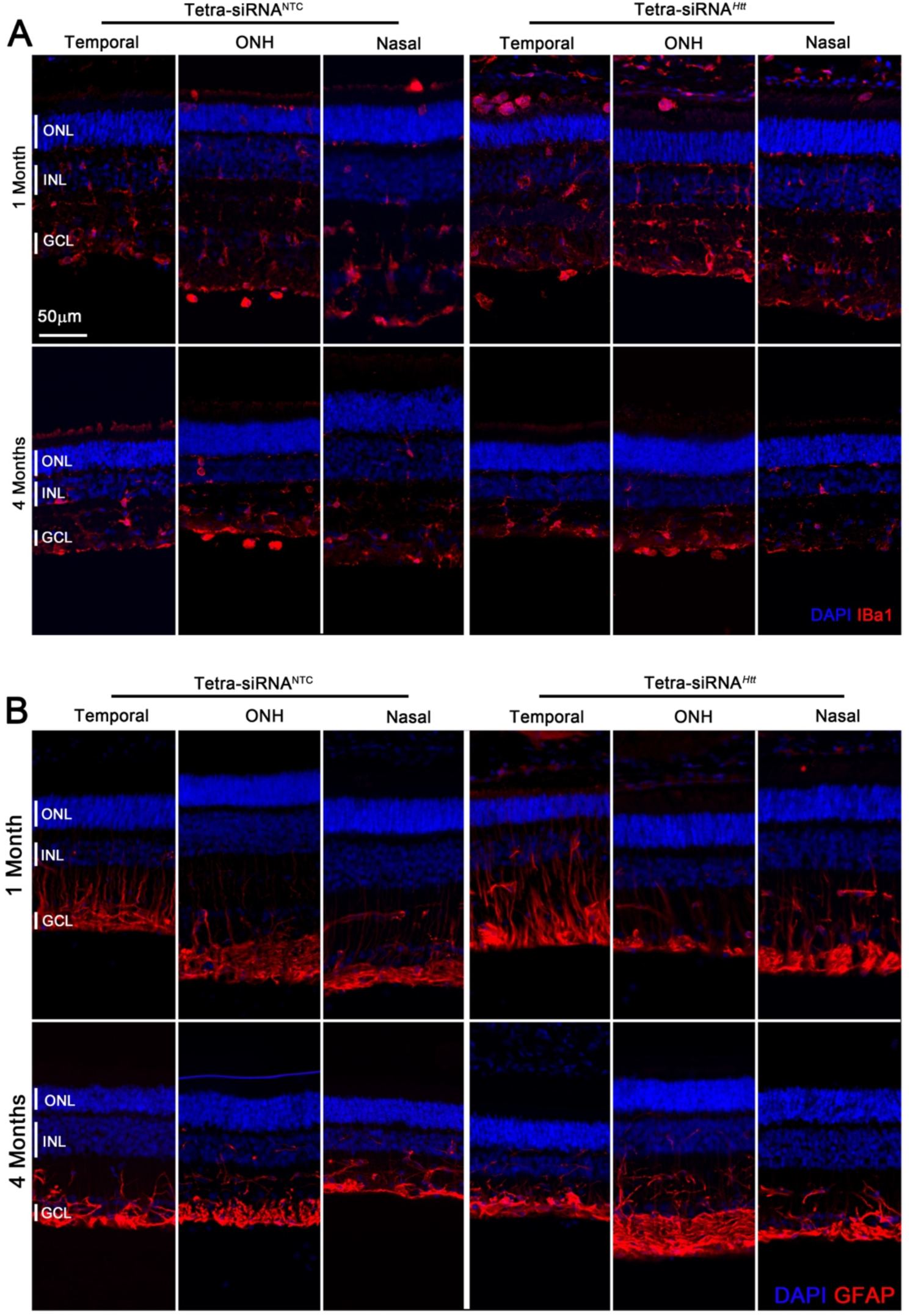
Safety of HTT silencing in porcine retina. (A and B) Retinal cross-sections stained for Iba1 (A: red signal) or GFAP (B: red signal) at 1-, and 4-months post intravitreal injection with 300μg of siRNA. The temporal side, where siRNA was injected shows slightly higher microglia activity and GFAP upregulation at 1-month post-injection. By 4-months post-injection microglia activity and GFAP expression appear normal again. Blue: nuclei marked with DAPI; red: Iba1 signal in (A) and GFAP signal in (B); ONL: outer nuclear layer; INL: inner nuclear layer; GCL: ganglion cell layer. Scale bar: 50μm. Images are representative images from 3 pigs per time point and siRNA.

